## Supporting Information for "*Yersinia* actively downregulates type III secretion and adhesion at higher cell densities"

for

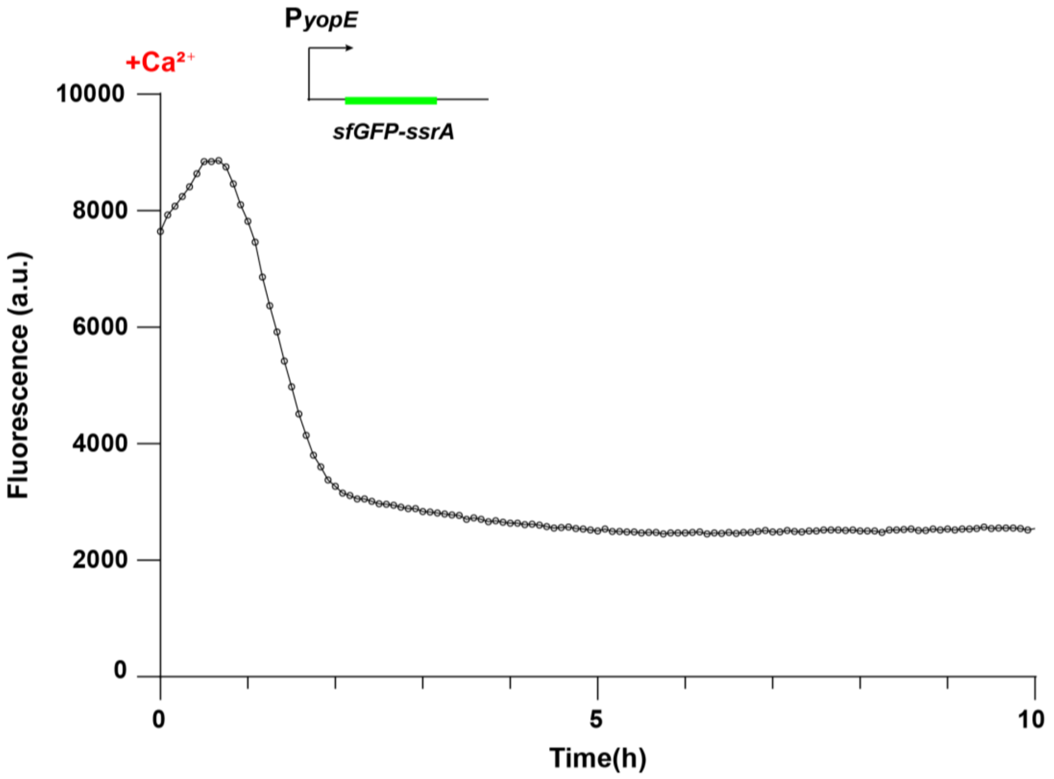


**Suppl. Fig. 1 – Kinetics of the P$_{yopE}$::*sfGFP-ssrA* activity assay.**

150 min after the induction of T3SS assembly by temperature shift to 37° in secreting conditions, T3SS activation in a wild-type strain carrying the P$_{yopE}$::*sfGFP-ssrA* reporter on the virulence plasmid was inhibited by adding 10 mM $CaCl_2$ to the medium (t=0). From t=0, fluorescence was measured over time. *n*=3, graph shows representative result.

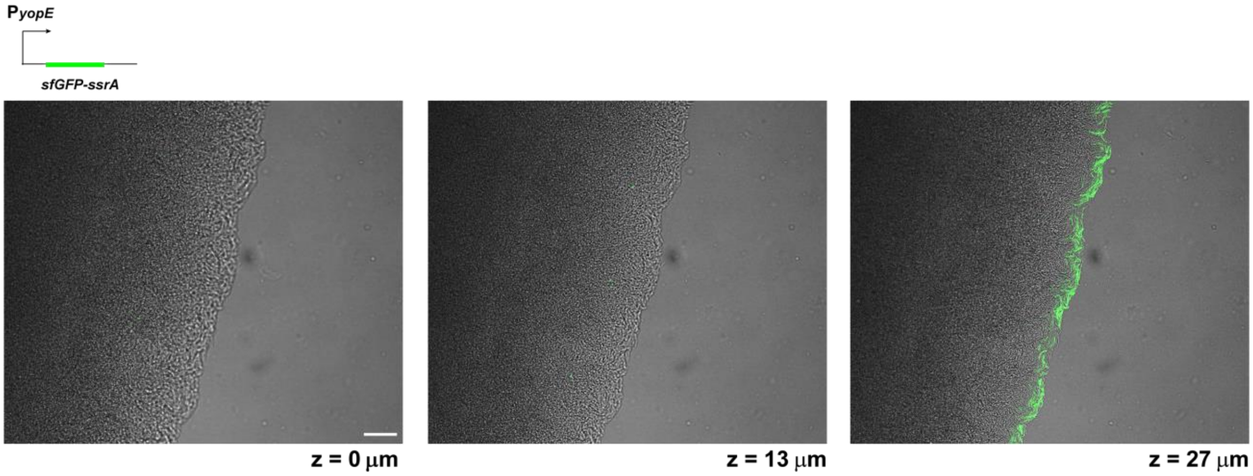


**Suppl. Fig. 2 – P$_{yopE}$::*sfGFP-ssrA* expression in *Y. enterocolitica* microcolonies.**

Image of confocal microscopy sections (z=0 µm, 13 µm, 27 µm) of a *Y. enterocolitica* Δ*sctW* P$_{yopE}$-*sfGFP-ssrA* microcolony section at 37°. T3SS activity, which results in a strong upregulation of the *yopE* promoter and cellular fluorescence, is only detected at the edge of the microcolony. Scale bar, 50 µm; *n*=3.

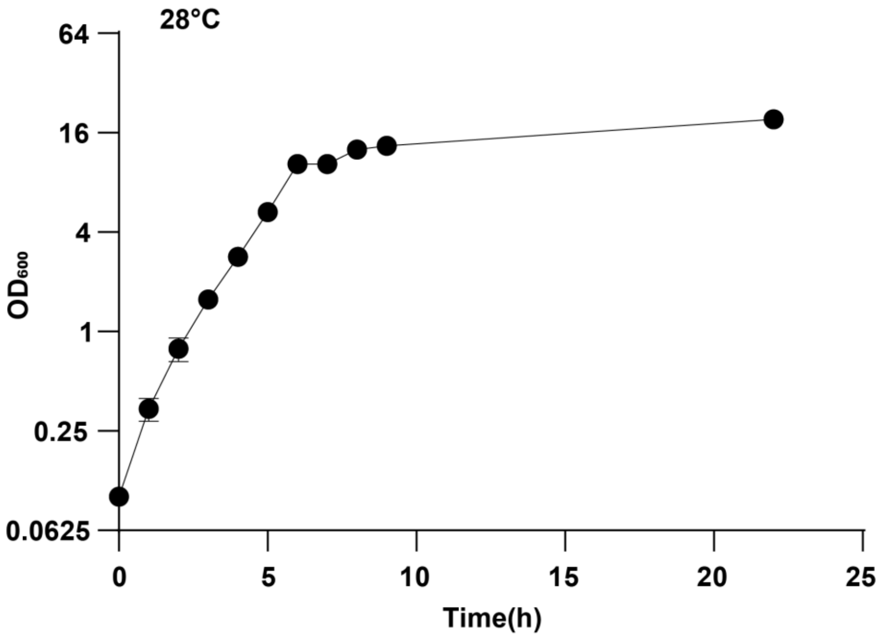


**Suppl. Fig. 3 – *Y. enterocolitica* reference growth curve.**

Optical density at 600 nm ($OD_{600}$) of *Y. enterocolitica* wild-type cultures (MRS40) incubated at 28°C. *n*=3; whiskers denote standard deviation.

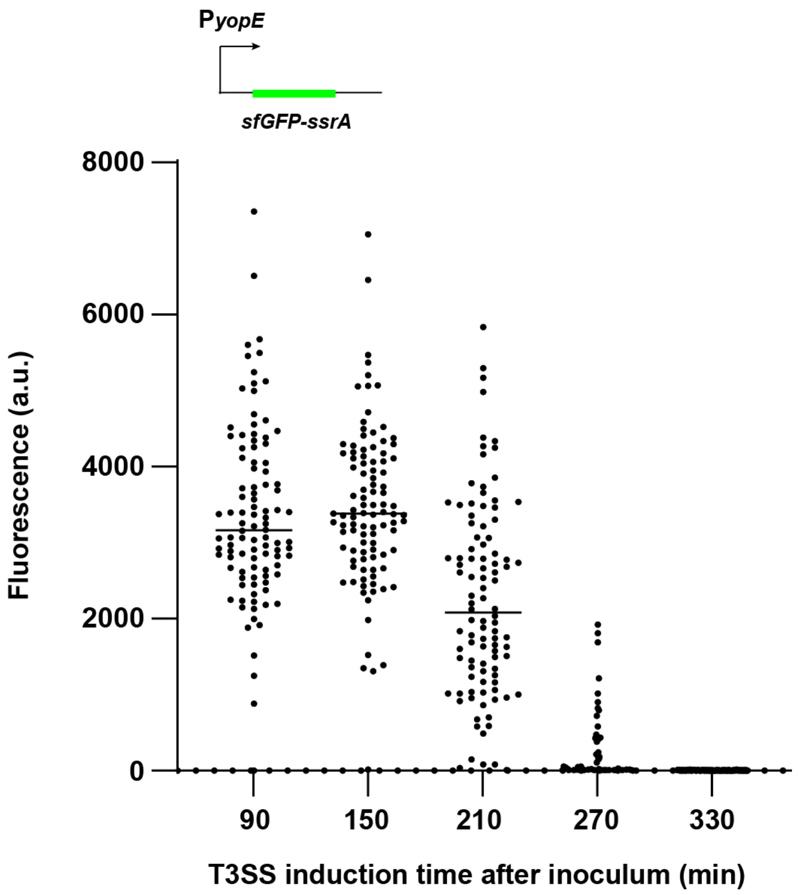


**Suppl. Fig. 4 – T3SS activity is repressed over time in growing cultures.**

T3SS reporter assay ($P_{yopE}$-*sfGFP-SsrA*) of *Yersinia* cells inoculated at $OD_{in}$=0.1 in secreting medium. Culture aliquots were shifted to 37°C to induce assembly of the T3SS at different time points post-inoculum as indicated. T3SS activity was measured at a single-cell level 2.5 h post-induction. *n*=3 independent experiments; each dot represents a single-cell measurement and the black bar represents the average of the fluorescence intensity.

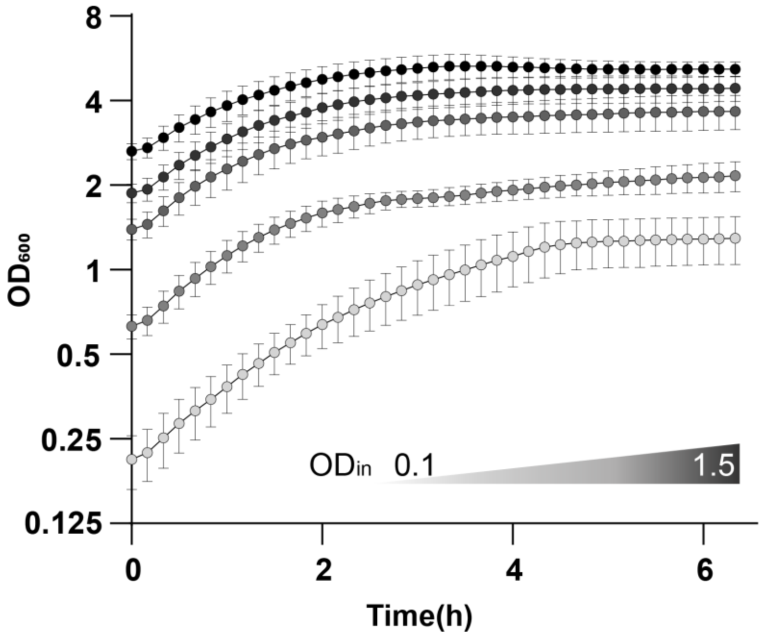


**Suppl. Fig. 5 – *Y. enterocolitica* growth curves at different $OD_{in}$.**

$OD_{600}$ of wild-type strain expressing $P_{yopE}$*::sfGFP-ssrA* under secreting conditions at the different $OD_{in}$ used in Fig. 2b (0.1, 0.3, 0.7, 1.0, 1.5). $OD_{600}$ was measured from the time of the shift to 37° (t=0). *n*=3, whiskers denote standard deviation.

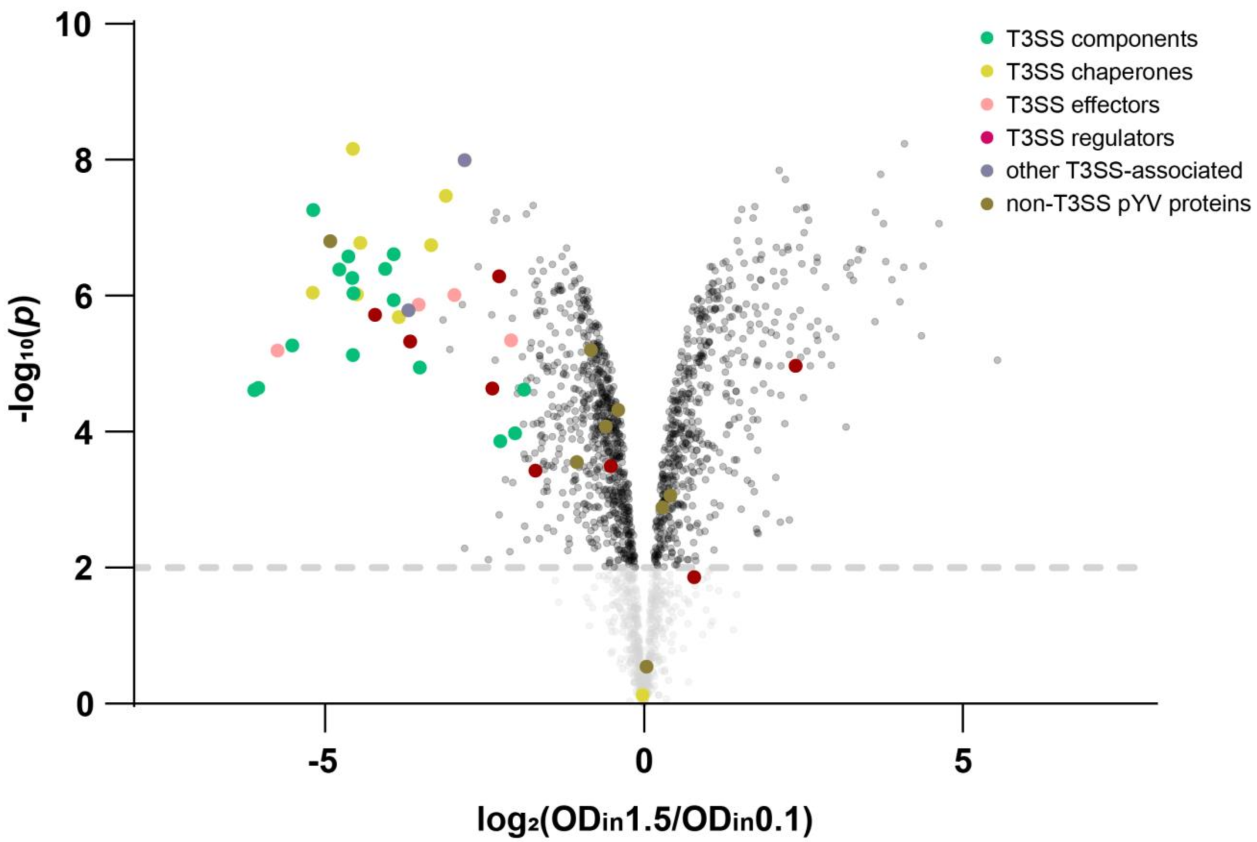


**Suppl. Fig. 6 – All T3SS gene categories are similarly downregulated at higher cell densities.**

Volcano plot showing differences in the expression of *Y. enterocolitica* proteins between cultures grown under secreting conditions at $OD_{in}$ 1.5 and 0.1. All proteins with at least three detected peptides are displayed. Representation of the plot shown in Fig. 3a (Table 1, Suppl. Table 2), highlighting different classes of proteins, colored according to their function.

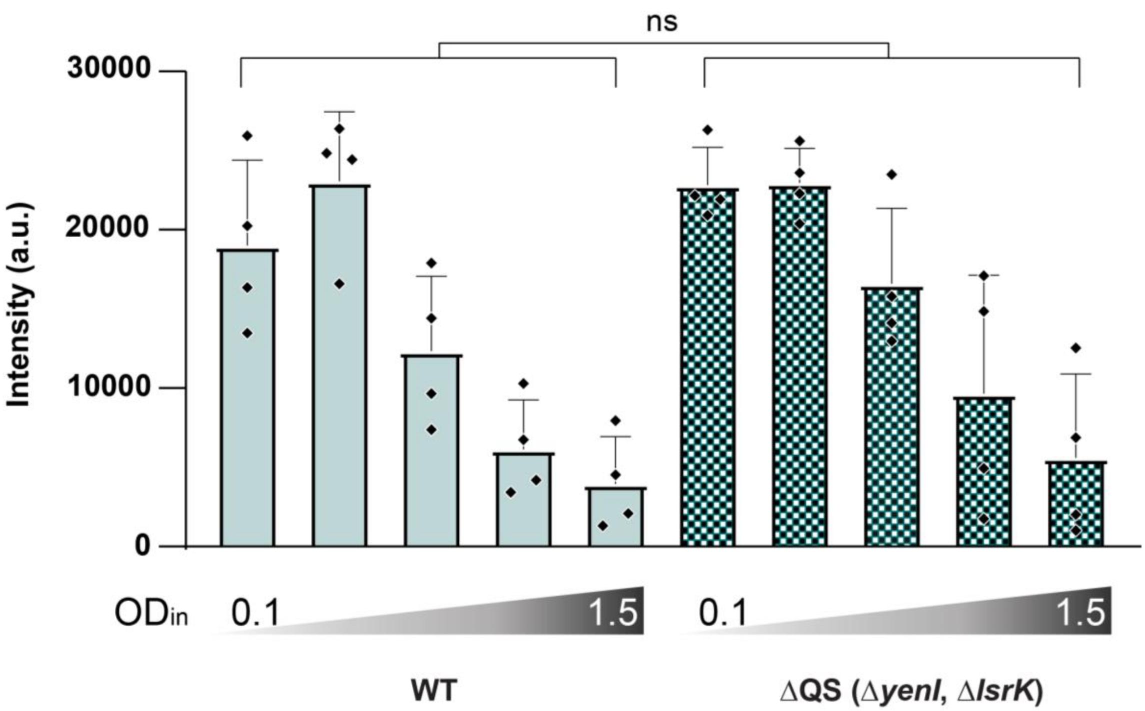


**Suppl. Fig. 7 – The density-dependent downregulation of the T3SS is not conferred by quorum sensing.**

Gel band intensity quantification of secretion assay of wild-type (WT, light blue) and quorum sensing deletion strain (ΔQS = *ΔyenI, ΔlsrK*, dotted bars) at $OD_{in}$ = 0.1, 0.3, 0.7, 1.0, 1.5 shown in Fig.4a. *n*=4, each spot represents a single measurement, whiskers denote standard deviation. For statistics, WT and ΔQS measurements were compared for each $OD_{in}$ using an unpaired t-test. Each comparison resulted in a statistically non-significant difference (ns, *p*>0.05), represented in the graph as grouped result.

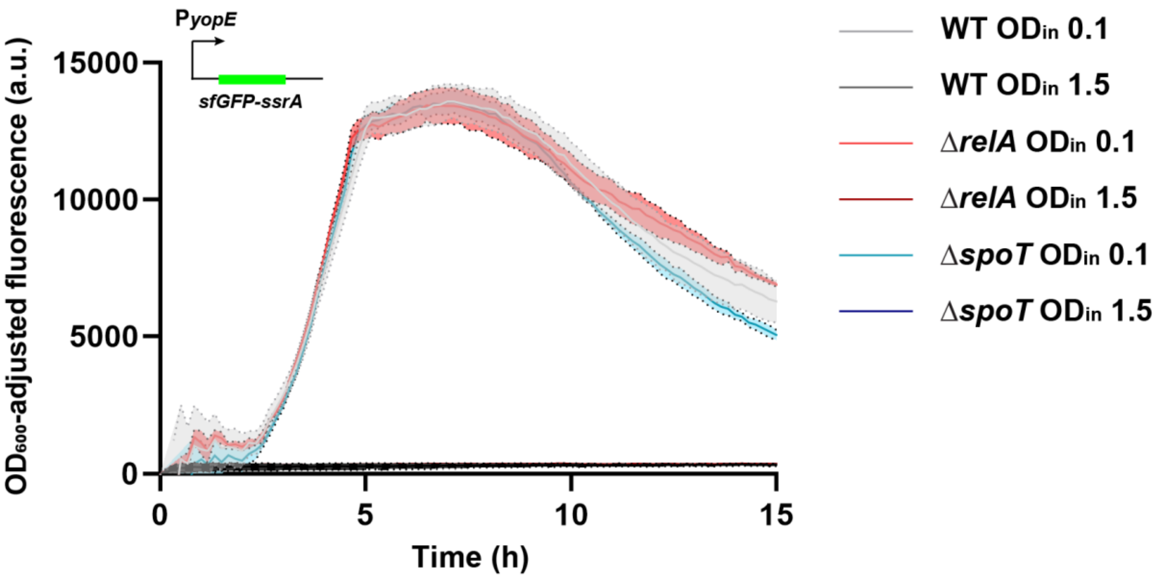


**Suppl. Fig. 8 – The stringent response regulators RelA and SpoT do not significantly contribute to the density-dependent downregulation of the T3SS.**

T3SS reporter assay ($P_{yopE}$-*sfGFP-SsrA*) of *Yersinia* cells at $OD_{in}$=0.1 and 1.5 in the indicated strains after shifting the culture to 37°C (t=0), which induces the expression of the T3SS. *n*=3, shadowed area denotes standard deviation.

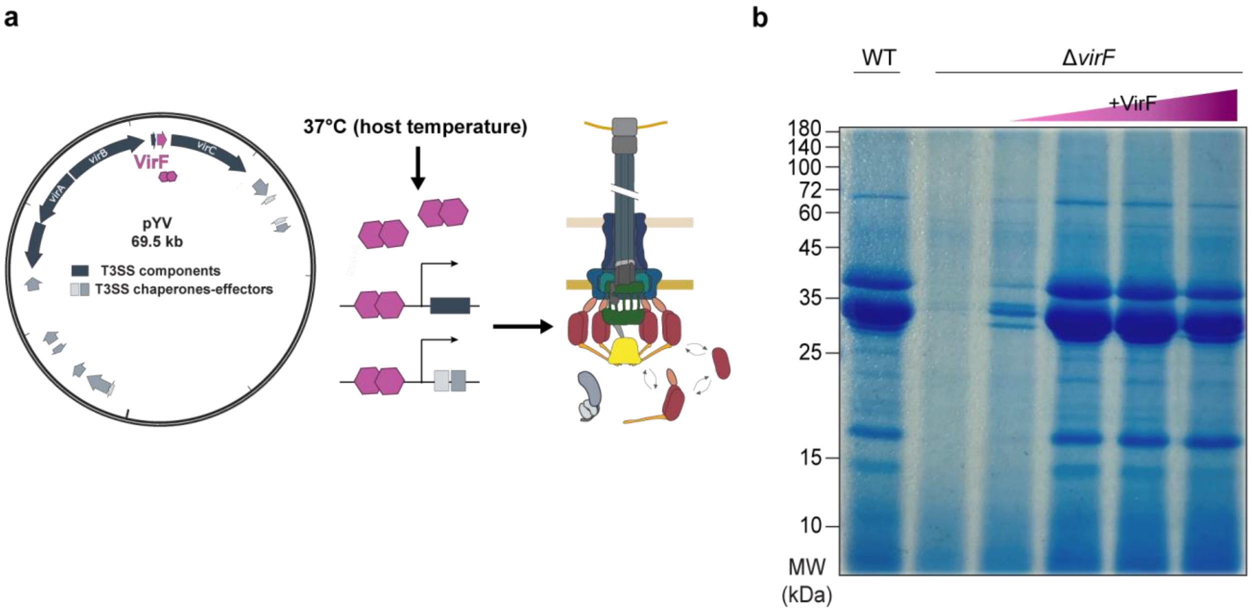


**Suppl. Fig. 9 – VirF is essential for T3SS secretion.**

**a)** Schematic display of the genes encoding for T3SS structural components (operons *virA*, *virB*, *virC*, and translocator operon (transl.), dark grey), T3SS effectors and chaperones (lighter shades of grey), and VirF (purple) on the pYV virulence plasmid (left), and the principle working mechanism of VirF (right). **b)** Secretion assay displaying the effectors secreted by the T3SS at $OD_{in} = 0.1$ in wild-type (WT) and Δ*virF* background. Lanes 2-6, increasing expression of VirF *in trans* (arabinose concentrations of 0, 0.001, 0.003, 0.008%, respectively). *n*=3, gel shows representative result.

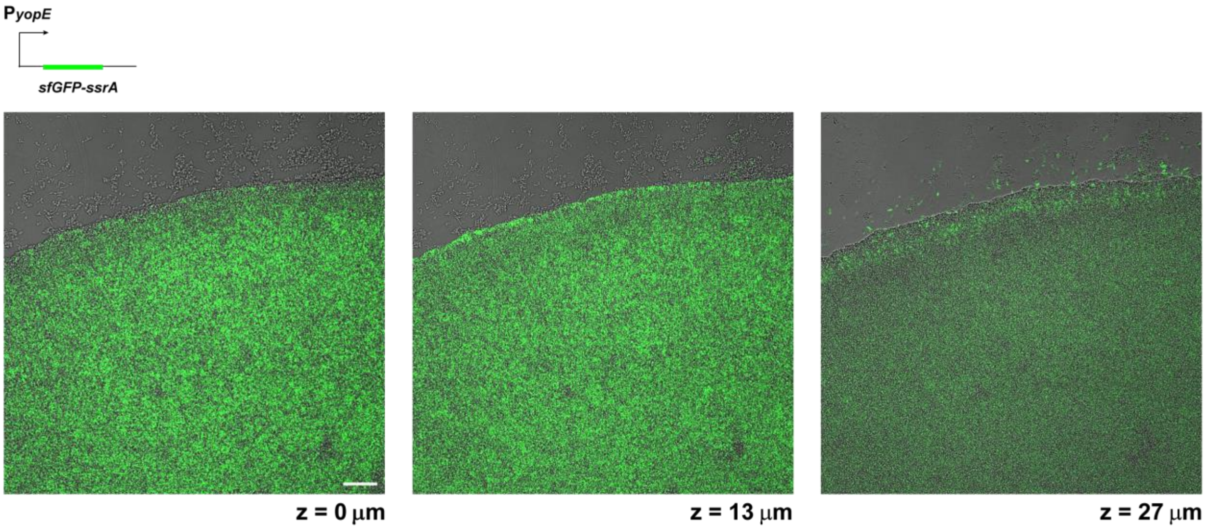


**Suppl. Fig. 10 – Additional expression of the transcriptional regulator VirF restores T3SS secretion in a *Yersinia* microcolony.**

Image of confocal microscopy sections (z=0 µm, 13 µm, 27 µm) of a *Y. enterocolitica* Δ*sctW* $P_{yopE}$-*sfGFP-ssrA* microcolony additionally expressing VirF (induced by 0.2% arabinose) at 37°C. T3SS activity, visualized by the sfGFP signal, is detected throughout the colony. Scale bar, 50 µm, *n*=3.

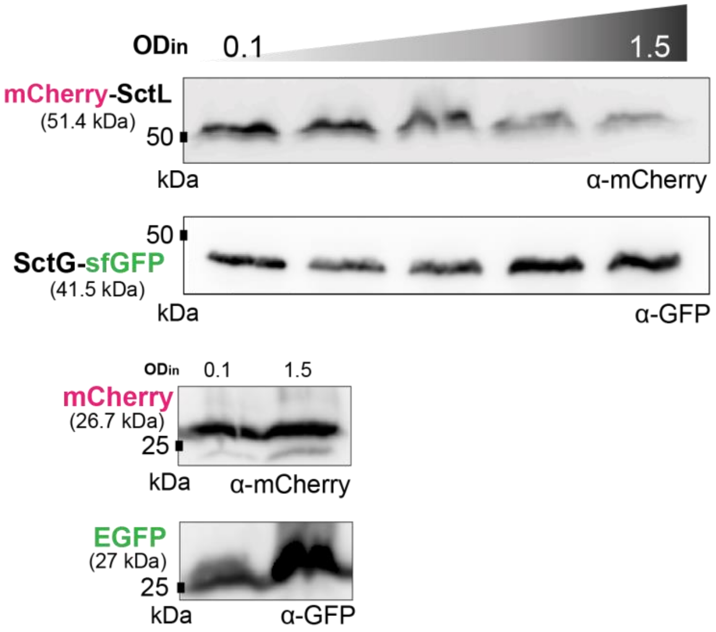


**Suppl. Fig. 11 – SctG protein levels are stable at higher local cell densities.**

Expression of mCherry-SctL (expected molecular weight 51.4 kDa) and SctG-sfGFP (expected molecular weight 41.5 kDa) expressed from the native genetic location in *Y. enterocolitica* in secreting medium at the $OD_{in}$ indicated. Immunoblot using antibodies directed against mCherry and EGFP as indicated. Control: Expression of mCherry (expected size 26.7 kDa) and GFP (expected size 27.0 kDa) from plasmid. *n*=3, blots show representative results.

```
CsrB Y. enterocolitica      GCCCGATAGGATCTGGC-GGAAAGGACGGTATCAGGATGGTGCCACTTCAGGATGAAGGA      59
CsrB Y. pseudotuberculosis  GCTGGATAGGATCTGGCGGAGAGGGAACGCATCTGGAAGATGTGTCTTCAGGACGAAGAA      60
                            **  ************* *  * ***  * *** *** * **   ******** **** *

CsrB Y. enterocolitica      CTCAGGGACTGCTTAGGATGAGTGAAGGGATGTTTCAGGAAGAAACAAAGGACACCTCCA      119
CsrB Y. pseudotuberculosis  CACAGGGACTGCTTAGGACGAGTGAAGGGACGTTTCAGGATGAAACAAGGGACACCTCCA      120
                            * **************** *********** ********* ******* ***********

CsrB Y. enterocolitica      GGATGGAGATTGAGAGCCAGTTCAGGATGATTGGTGGGTTAGGATAACTTCAGGATTGGC      179
CsrB Y. pseudotuberculosis  GGATGGAGATTGAGAGCCAGTTCAGGATGATTGGTGGGTTAGGATAGCCTAAGGATTAAC      180
                            ********************************************** * * ******  *

CsrB Y. enterocolitica      ACTGGGATGGTGTAGGACAACGCGACGGATTGCTGGTTAGGATAACCGCACGGAAAAGTT      239
CsrB Y. pseudotuberculosis  GCCGGGATGGTGTATAACATTGCGATGGATTGCTGGTTAGTATAACCATACGGAAAAGTT      240
                             * ***********  ***  **** ************** ******  ***********

CsrB Y. enterocolitica      TTCAAGGATTGAGCAGGGAGCATCACTTTTAGCTGGATTGCTATGAAACGAATAGAGGGG      299
CsrB Y. pseudotuberculosis  TTCAGGGATTGAGCAGGGAGCATCAATTTTAGCTGGATTGCTATAAAACGAATTGAGGGG      300
                            **** ******************** ****************** ******** ******

CsrB Y. enterocolitica      TACTGGTAAAACAGTACCCCTTTT  323
CsrB Y. pseudotuberculosis  TACTGGTAAAACAGTACCCCTTTT  324
                            ************************


CsrC Y. enterocolitica      TGACTATTTTTTGTAACTCATGGTTTTTTAACAGTGCGGGATGTACTGGC---AAGGAGC      57
CsrC Y. pseudotuberculosis  ----------------------ATACAAGGAATGTAATGGATGTACGGGAGCCAAGGACG      38
                                                   *      *  **   ******** **    *****

CsrC Y. enterocolitica      GTAATCACTTAGGAAGAGTGGGGTATGCTTAAGGAATGTAATGGATATACTGTGAGCCAG      117
CsrC Y. pseudotuberculosis  CTATAAACGGAG-----CTAATGTAAAGGGATGTTAGGACACTGGCCGGAGCGCCGGGCG      93
                             **   **  **      *   ***     * *  * *  *  *           *   *

CsrC Y. enterocolitica      GGACACCTTCAGGGTTGGGGGGATGGCAAGGATGGCGAATTGCAGTAGGGAGAAACCGGG      177
CsrC Y. pseudotuberculosis  AATCGCCTTCAGGGTTTGAGGGGTGGCAAGGATCGCGTATTGCAGGAGGGAGAAACCGGG      153
                               * *********** * *** ********** *** ******* **************

CsrC Y. enterocolitica      ACGTTATATCAGTGTAGGGAATGCACAATAAGGATATCCTTCCGCAGAAGGTGCGAAAAA      237
CsrC Y. pseudotuberculosis  ACGTTATCTAAGTGCAGGGAGTGCGCTGTAAGGATATCCTTCCGCAGAAGGTGCGAAAAA      213
                            ******* * **** ***** *** *  ********************************

CsrC Y. enterocolitica      AGGCGACAGGTTAATCTGCCGCCTTTTTTCTTTCTCTTCTT   278
CsrC Y. pseudotuberculosis  AGGCGACAGGTTAACCTGCCGCCTTTTTTCTTTCTCTTCTT   254
                            ************** **************************
```

**Suppl. Fig. 12 – Sequence conservation of CsrB and CsrC in *Y. enterocolitica* and *Y. pseudo-tuberculosis*.**

DNA sequence alignment using Clustal Omega. Nucleotides that are identical in both sequences are indicated by asterisks.

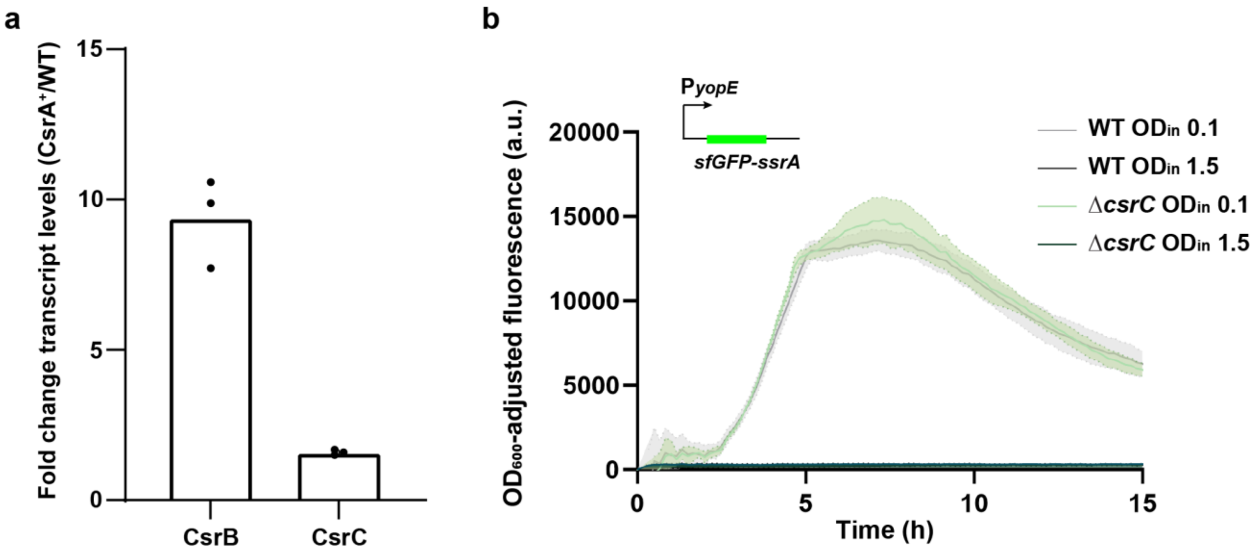


**Suppl. Fig. 13 – Manipulations of the CsrABC system are efficiently compensated.**

**a)** Changes in the transcript levels of CsrB and CsrC at $OD_{in}$=1.5 upon CsrA overexpression ($CsrA^{+}$, 0.2% arabinose) in comparison to the wildtype strain (WT), as measured by reverse transcription quantitative PCR. Additional CsrA expression is compensated by higher levels of CsrB and, to a lower degree, CsrC. **b)** T3SS reporter assay ($P_{yopE}$-*sfGFP-ssrA*) at $OD_{in}$ 0.1 and 1.5 in the indicated strains after shifting the culture to 37°C (t=0), which induces the expression of the T3SS. *n*=3 for all panels; shadowed area in b) denotes standard deviation,.

| $OD_{in}$ | 0.1 | 0.3 | 0.7 | 1.0 | 1.5 |
|---|---|---|---|---|---|
| # bacteria | 369 | 369 | 369 | 369 | 369 |
| 0 foci | 39 | 131 | 288 | 350 | 364 |
| 1 focus | 92 | 103 | 59 | 16 | 5 |
| 2 foci | 70 | 59 | 16 | 1 | 0 |
| 3 foci | 76 | 45 | 5 | 1 | 0 |
| 4 foci | 61 | 19 | 1 | 0 | 0 |
| 5 foci | 20 | 6 | 0 | 0 | 0 |
| 6 foci | 8 | 5 | 0 | 0 | 0 |
| 7 foci | 3 | 1 | 0 | 0 | 0 |
| ≥8 foci | 0 | 0 | 0 | 1 | 0 |

**Suppl. Table 1 – Quantification of EGFP-SctQ foci per bacterium at different bacterial densities.**

Quantification of EGFP-SctQ foci per bacterium, corresponding to assembled injectisomes, visualized by fluorescence microscopy at the indicated $OD_{in}$ values. Corresponds to Fig. 2c.

| Protein | $Log_2$ intensity ratio | *p* value | Individual replicate $log_2$ intensity values $OD_{in}$ = 0.1 | | | $OD_{in}$ = 1.5 | | | # pept. |
|---|---|---|---|---|---|---|---|---|---|
| Adhesin YadA | -4.94 | 1.6E-07 | 33.42 | 33.44 | 33.51 | 28.23 | 28.61 | 28.72 | 65 |
| Resolvase TnpR | -1.26 | 3.6E-04 | 24.11 | 24.09 | 24.14 | 22.50 | 23.04 | 23.00 | 6 |
| Antitoxin ParD | -0.83 | 6.3E-06 | 28.76 | 28.79 | 28.83 | 27.98 | 28.01 | 27.90 | 14 |
| Toxin ParE | -0.60 | 8.3E-05 | 25.93 | 25.88 | 25.93 | 25.37 | 25.22 | 25.36 | 4 |
| Transposase TnpA | -0.43 | 2.8E-03 | 22.55 | 22.25 | 22.41 | 21.97 | 21.96 | 21.99 | 2 |
| Plasmid partitioning protein SpyA | -0.40 | 4.8E-05 | 28.32 | 28.33 | 28.35 | 27.94 | 27.92 | 27.94 | 21 |
| Plasmid partitioning protein SpyB | 0.04 | 2.8E-01 | 28.43 | 28.39 | 28.38 | 28.46 | 28.46 | 28.42 | 20 |
| Arsenate reductase ArsC | 0.29 | 1.3E-03 | 26.37 | 26.38 | 26.29 | 26.66 | 26.65 | 26.58 | 14 |
| Arsenic resistance protein ArsH | 0.42 | 8.7E-04 | 25.51 | 25.34 | 25.40 | 25.77 | 25.89 | 25.83 | 10 |
| Arsenite inducible repressor ArsR | 0.79 | 6.9E-03 | 20.40 | 20.89 | 21.07 | 21.49 | 21.71 | 21.55 | 3 |

**Suppl. Table 2 – Density-dependent regulation of expression of non-T3SS components encoded on the *Yersinia* virulence plasmid.**

While the adhesin YadA is downregulated similarly to the T3SS components (Table 1), most non-T3SS proteins encoded on the *Yersinia* pYV virulence plasmid are not strongly affected by the density-dependent downregulation. Label-free quantitative mass spectrometry in the total proteome of a ΔHOPEMTasd wild-type strain at the different growth conditions indicated, experiment and display format as shown in Table 1.

| Protein *(gene name)* | Log₂ intensity ratios | | Individual replicate $log_2$ intensity values | | | | | | | | | # pept. |
|---|---|---|---|---|---|---|---|---|---|---|---|---|
| | $OD_{in}$ 1.5 / $OD_{in}$ 0.1 | Stat. / $OD_{in}$ 1.5 | $OD_{in}$ = 0.1 | | | $OD_{in}$ = 1.5 | | | Stationary | | | |
| Osmotically-inducible protein Y *(osmY)* | 0.77 | 1.63 | 26.23 | 26.17 | 26.22 | 26.92 | 27.00 | 27.03 | 28.65 | 28.55 | 28.65 | 16 |
| Acetate operon repressor *(iclR)* | -0.29 | 2.02 | 23.66 | 23.41 | 23.60 | 23.29 | 23.31 | 23.22 | 25.19 | 25.30 | 25.39 | 16 |
| Periplasmic protein CpxP *(cpxP)* | -0.21 | 2.15 | 28.88 | 28.81 | 28.63 | 28.60 | 28.56 | 28.53 | 30.57 | 30.62 | 30.92 | 38 |

**Suppl. Table 3 – The density-dependent downregulation of the T3SS differs from stationary phase response.**

Label-free quantitative mass spectrometry of the known stationary response proteins CpxP, IclR and OsmY [1–4] in the total proteome of different growth conditions, as indicated. Stat., stationary cultures. # pept., number of total detected peptides. Display format as shown in Table 1 and Suppl. Table 2.

| Protein | $Log_2$ intensity ratio | *p* value | Individual replicate $log_2$ intensity values | | | | | | # pept. |
|---|---|---|---|---|---|---|---|---|---|
| | | | $OD_{in}$ = 0.1 | | | $OD_{in}$ = 1.5 | | | |
| CsrA | 0.52 | 0.17 | 23.82 | 24.56 | 25.06 | 24.83 | 25.01 | 25.15 | 7 |

**Suppl. Table 4 – CsrA levels are not not significantly altered at different densities.**

Label-free quantitative mass spectrometry quantification of CsrA in the total proteome of a ΔHOPEMTasd wild-type strain at the different growth conditions indicated, experiment and display format as shown in Table 1.

| Strain | Genotype | Reference |
|---|---|---|
| MRS40 | Wild-type *Y. enterocolitica* E40 *ΔblaA* | [5] |
| IML421*asd* (ΔHOPEMTasd) | MRS40 *yopH*$_{Δ1\text{-}352}$ *yopO*$_{Δ65\text{-}558}$ *yopP*$_{43}$ *yopE*$_{5}$ *yopM*$_{18}$ *yopT*$_{135}$ *Δasd* | [6] |
| AD4085 | IML421*asd egfp-sctQ* | [6] |
| AD4615 | MRS40 $P_{yopE}$*::sfGFP-ssrA* | This study |
| CH4010 | MRS40 *mCherry-sctL sctG-sfgfp* | [7] |
| FE005 | IML421*asd* $P_{yopE}$*::sfGFP-SsrA ΔyadA* | This study |
| FE014 | IML421*asd ΔvirF* | [8] |
| FE015 | MRS40 *ΔrpoS* | This study |
| FE022 | IML421*asd ΔyenI ΔlsrK* | This study |
| FE027 | $P_{yopE}$*::sfGFP-SsrA ΔcsrC* | This study |
| FE028 | $P_{yopE}$*::sfGFP-SsrA ΔrelA* | This study |
| FE029 | $P_{yopE}$*::sfGFP-SsrA ΔspoT* | This study |

| Plasmids | Genotype | Reference |
|---|---|---|
| pAD477 | pBAD::mCherry (expression plasmid for mCherry expression) | [9] |
| pAD492 | pBAD::EGFP (expression plasmid for EGFP expression) | [7] |
| pAD716 | pKNG101-PyopE-sfGFP-SsrA (mutator for P*yopE* reporter system) | This study |
| pBAD-His B | pBR322-derived expression vector | Invitrogen |
| pFE002 | pKNG101-*ΔrpoS* (mutator for deletion of the *rpoS* gene) | This study |
| pFE003 | pBAD::*rpoS* (expression plasmid for RpoS expression) | This study |
| pFE010 | pBAD::*virF* (expression plasmid for VirF expression) | [8] |
| pFE018 | pKNG101-*ΔyenI* (mutator for deletion of the *yenI* gene) | This study |
| pFE021 | pKNG101-*ΔlsrK* (mutator for deletion of the *lsrK* gene) | This study |
| pFE022 | pBAD::*csrA* (expression plasmid for CsrA overexpression) | This study |
| pFE025 | pKNG101-*ΔcsrC* (mutator for disruption of the *csrC* gene) | This study |
| pFE026 | pKNG101-*ΔrelA* (mutator for disruption of the *relA* gene) | This study |
| pFE027 | pKNG101-*ΔspoT* (mutator for disruption of the *spoT* gene) | This study |
| pLJM31 | pKNG101-*ΔyadA* (mutator for disruption of the *yadA* gene) | [10] |
| pKNG101 | *oriR6K sacBR*$^{+}$ *oriTRK2 strAB*$^{+}$ (suicide vector for homologous recombination) | [11] |

**Suppl. Table 5 – Strains and plasmids used in this study**

| Primer name | Sequence (5´→ 3´) | Used for |
|---|---|---|
| AD1117 | AAGGTCTCGGGCCCCGATAACCGGTTCAATAGTATCTGG | pAD716 |
| AD1118 | GTCCTTGTAGTCACCTCCCAATTGAAAGATCTTATTTTCATGACTATTTATTCCCTTGGCT | pAD716 |
| AD1119 | CAATTGGGAGGTGACTACAAGGACGACGATGATAAGTGATATGGATAAAAACAAGGGGGTAG | pAD716 |
| AD1120 | AAGGTCTCTCTAGAGATTTGCTCTGACATGCGCC | pAD716 |
| AD1498 | TATAGGTCTCGGGCCCTGAATCGAAAGTAACTTGCGGTTGGTAG | pFE002 |
| AD1499 | TATAGGTCTCTCTAGATTACTCGGTTCATGATCTAATTTATGA | pFE002 |
| AD1500 | TATACCATGGGTAGCCAAAATACGCTGAAAGTTAACGAGT | pFE003 |
| AD1501 | TATAAAGCTTTTCGCGGAACAATGCTTCGATGCTCAGG | pFE003 |
| AD1835 | TATAAAGCTTGCCTGTGGTTGCTATTTTAGTAAGAC | pFE010 |
| AD1856 | TATAGGTCTCCCATGGGTGCATCACTAGAGATTATTAAATTAGAATGGGC | pFE010 |
| AD1901 | TATAGGTCTCGGGCCCTATCTCCAATGGAAGCGACGATAGTATC | pFE018 |
| AD1902 | TTTTTATTTTAGCTTCAACTCAATGCCAAGC | pFE018 |
| AD1903 | GAAGCTAAAATAAAAACCAAAGTTATTTAATTAAAACAACATCGT | pFE018 |
| AD1904 | AAGGTCTCTCTAGAGACGAGAAAAACAAATTTTTTATTAGTAAAAAGTGG | pFE018 |
| AD1940 | TATAGGTCTCGGGCCCAGGATGCGCCATCATCAGCAACG | pFE021 |
| AD1941 | GATCGAACAAAAATTAGCTATCTTACTGGCCAGCT | pFE021 |
| AD1942 | GCTAATTTTTGTTCGATCCTCAGCTTCAATGTG | pFE021 |
| AD1943 | AAGGTCTCTCTAGACTGATGAGTTGACCGCAAGATCA | pFE021 |
| AD2232 | TATAGGTCTCGGGCCCCTGAAGGTGACTCACCCAG | pFE025 |
| AD2233 | TATAGGTCTCTCTAGAGATGTACTGGCAAGGAGCG | pFE025 |
| AD2234 | TATAGGTCTCGGGCCCACATATACCCGATCGTCAAATACC | pFE026 |
| AD2235 | TATAGGTCTCTCTAGAGCAGATGAAGGGATCAAAGCC | pFE026 |
| AD2236 | TATAGGTCTCGGGCCCTAAAGAAGTCGATACGTGCTATCG | pFE027 |
| AD2237 | TATAGGTCTCTCTAGAAAAGTGGATAGGGCTGGCG | pFE027 |
| AD2032 | TATACCATGGGTCTTATTCTGACTCGTCGAGTTGGT | pFE022 |
| AD2033 | TATAAAGCTTCAGTAAGTCGTCGGTTGAGACT | pFE022 |
| AD1977 | TCACCAACAACATTCCACAG | qPCR *gyrB* |
| AD1978 | TTCGACCGCAGTTTTTACC | qPCR *gyrB* |
| AD1981 | CCATTATCTCGCACAGCAC | qPCR *sctG* |
| AD1982 | TAAAGTTCACCTTCCCCGC | qPCR *sctG* |
| AD1983 | AGACGAGACAATGCCACAC | qPCR *virF* |
| AD1984 | GCAGAGCCGAGAGGAATAAAG | qPCR *virF* |
| AD2037 | GACGGTATCAGGATGGTGCC | qPCR *csrB* |
| AD2038 | TCCATCCTGGAGGTGTCCTT | qPCR *csrB* |
| AD2039 | TCCGGCCTGTGTCCATAAAC | qPCR *csrC* |
| AD2040 | AACAGTGCGGGATGTACTGG | qPCR *csrC* |

**Suppl. Table 6 – Oligonucleotides used in this study**

Nucleotide sequence of oligonucleotides used for the construction of plasmids or quantitative PCR (qPCR).
